# Method for Zero-Waste Circular Economy using worms for plastic agriculture: Augmenting polystyrene consumption and plant growth

**DOI:** 10.1101/2020.05.29.123521

**Authors:** Samuel Ken-En Gan, Ser-Xian Phua, Joshua Yi Yeo, Zealyn Shi-Lin Heng, Zhenxiang Xing

## Abstract

Polystyrene (PS) is one of the major plastics contributing to environmental pollution with its durability and resistance to natural biodegradation. Recent research showed that mealworms (*Tenebrio molitor*) and superworms (*Zophobas morio*) are naturally able to consume PS as a carbon food source and degrade them without observable toxic effects. In this study, we explored the effects of possible food waste contamination and use of worm frass as potential plant fertilizers. We found that small amounts of sucrose and bran increased PS consumption and that the worm frass alone could support dragon fruit cacti (*Hylocereus undatus*) growth, with superworm frass in particular, supporting better growth and rooting than mealworm frass and control media over a fortnight. As known fish and poultry feed, these finding present worms as a natural solution to simultaneously tackle both the global plastic problem and urban farming issue in a zero-waste sustainable bioremediation cycle.

## 1. Introduction

Styrofoam, or polystyrene (PS), are light polymers with low heat conductivity [1,2] that can be synthesized to different shapes and sizes, allowing it to be widely used world-wide. While ubiquitous, its resistance to degradation causes its waste to accumulate, leading to pollution [3]. Even though PS waste can be incinerated, this releases toxic fumes, causing air pollution [4–6], pushing for the search of better alternative PS waste management methods.

Within the darkling beetle (*Tenebrionidae*) family, superworms (*Zophobas morio*) and mealworms (*Tenebro molitor*) are naturally voracious agricultural insect pests. They are protein rich food sources [7], with mealworms recently approved to be safe for human consumption in the EU [8]. Mealworms were found to naturally consume, metabolize and mineralize the carbon in PS [9], an ability found to be conferred by the commensal gut bacteria [10]. As larvae, they can be bred at high density to excrete nitrogen [11] and chitin rich [12,13] frass waste that was shown capable of substituting traditional NPK (Nitrogen, Phosphorus, and Potassium) fertilizers [14–16] in a circular economy [15]. The use of insect frass as alternatives to commercial fertilisers was recently shown with black soldier fly frass demonstrating to support the growth of maize [17,18] and ryegrass [19].With reports that superworms were able to consume PS at a higher rate than mealworms [20], there is promise of superworms joining mealworms in the fight against PS waste.

Given that the bulk of PS waste are food contaminated packaging that need to be cleaned prior to many current recycling methods, the natural plastic degradation by worms seems to be a better solution, especially when considering that the worms are food sources, and that their frass are natural fertilizers to form a zero-waste (a goal as defined by the Zero Waste International Alliance [21] and Eco Cycle Solutions Hub [22] conversion of PS waste.

To evaluate the methodology of using augmenting the natural ability of these worms to consume PS waste, we performed a feasibility check on 1) the effect of food additives on PS consumption by mealworms and superworms; and 2) the use of PS-fed superworm and mealworm frass to support dragon fruit cacti (*Hylocereus undatus*, chosen as an easy growing indoor fruit plant) rooting and growth.

## 2. Materials and Methods

### 2.1. Insect rearing and frass collection

Superworms (*Zophobas morio*) and mealworms (*Tenebro molitor*) fed on bran were purchased from fish feed stores (Clementi, Singapore). They were weighed and placed in polypropylene (PP) containers (impervious to the worms) with the food condiments added to PS balls (see Figure 1A & B for experimental setup). Worm frass were sifted using a mesh sieve to remove uneaten PS/food and worm parts. The worms were kept in a constant humidity of ~50% and a temperature of ~25°C (previously reported to be ideal for PS consumption by worms [23]) with the environment monitored by assembled Arduino devices (not shown).

**Figure 1.**
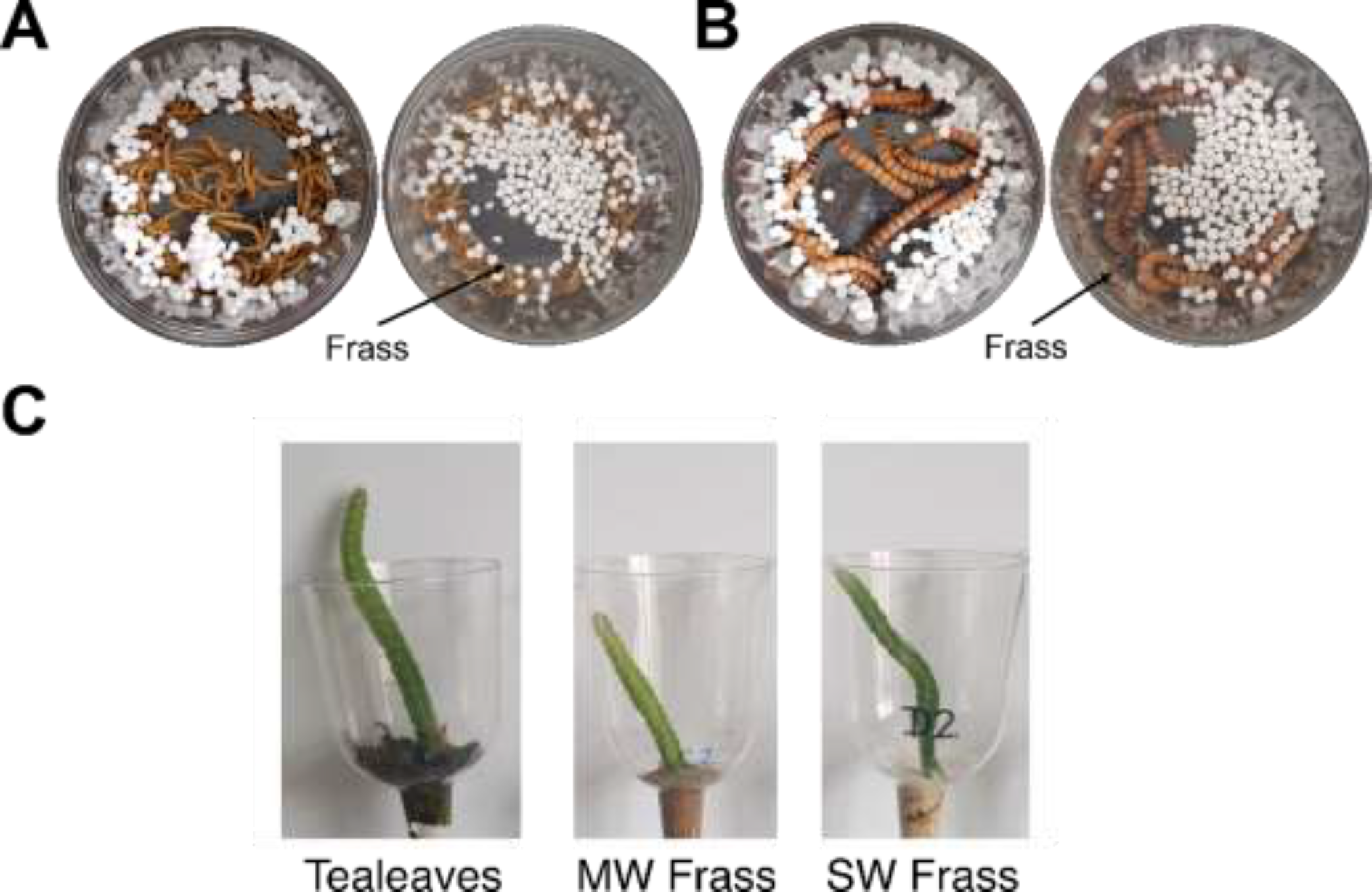
Representative images of the setups for testing PS consumption rates by (A) Mealworms (MW); (B) Superworms (SW). For both A and B, the left are the initial setups, and the right showed the setup after four days with frass produced from the PS consumption alone. (C) Setup of the dragon fruit cacti grafted onto the test media of tea leaves, MW, and SW frass in transparent plastic holders with a funnelled end to allow the grafted cacti to be covered by less frass while upright.

### 2.2. PS consumption rate experiments

The natural rate of PS consumption (mg of PS / g of worm per day) by multiple replicate setups of superworms and mealworms were measured in multiple setups of total worm weights between 6.22-10.76 g and 300-390 mg of PS balls (Figure 1A & B) with diameters between 0.4-0.5 cm (Art friend, Singapore). To test food contamination, PS balls were premixed with 25 mg of either cinnamon powder (Masterfoods, Australia, chosen for the strong smell), bran (Bob’s Redmill, America, the original supplied food source), table sucrose (Lippo group, given sugar is a common insect attractant) or no additive (control) carried out in sextuplicates using the same batch of additives. 0.9 ml of water was added to the mix of food additives and PS balls, and the unconsumed PS balls were collected after four days and weighed on an analytical balance. The final total live worm weights were used for calculation. Worm weight changes are shown in supplementary data (Table S1).

### 2.3. Worm frass and Dragon fruit cacti (Hylocereus undatus) experiment setups

For accurate analysis of frass, the worms in the experiments were reared solely on PS balls at least a week before renewed frass collection.

*Hylocereus undatus* cacti of the same original seed-grown pot (bought in a local supermarket), were grafted successfully multiple times on used Chinese tea leaves (termed tea leaves) as media for five years. Given previous reports of its benefits in plant growth [24,25], tea leaves were used as a positive control to study the effects of the worm frass. The worm frass experiments were setup using the same grafting method on the 33 selected cacti branches (3 media conditions × 11 cacti replicates) of similar size onto the test media and grown in transparent plastic pots (see Figure 1C for set-up) with funnelled bottoms to support upright growth. Used tea leaves, superworm frass, and mealworm frass covered the bottom of the grafted cacti, forming the soil line.

The grafted cacti were placed against a window ledge and watered twice a week to wet the media. The height of the straightened grafted cacti (end to end, excluding roots) were measured before grafting and after a fortnight. Only rooting that occurred below the soil line were considered given the existing presence of aerial roots for the cacti. Dead cacti (supplementary Table S2) were also recorded.

### 2.4. GC-MS analysis of superworm frass

Gas chromatography-mass spectrometry (GC-MS) was used to analyse the worm frass and worms for the presence of PS and its by-products. Superworms and mealworms were fed with bran or PS balls exclusively for two weeks. For analysis of PS and its by-products, the worms were starved for five days to ensure no undigested PS were left in the body and the worms killed for the analysis. PS balls, frass (20 mg), and worm carcasses were dissolved in dichloromethane and incubated in 2 ml microfuge tubes for 10 minutes and subsequently centrifuged (14.8k RPM, 5 minutes using table top centrifuge) to remove undissolved solids. The solvent soluble fraction were syringe filtered using 0.45 μm teflon filters and analysed using the HP 6890 gas chromatography HP-5MS column and HP 5973 mass spectrometry system. The GC oven temperature was set to 50°C for 1 minute and 250°C for 5 minutes at the ramp up rate of 10°C/minute

### 2.5. Statistical analysis

The mean, standard error and fold change were calculated and analysed using Microsoft Excel (version 15.0). Two-tailed unequal variances T-test was performed to compare the effect of food additives on superworm and mealworm polystyrene consumption, and the effect of different media on cacti height. For the comparison of the different media on cacti rooting frequency, Pearson’s Chi-squared test was performed. To compare differences in worm weight pre- and post-feeding on polystyrene, two-tailed paired T-test was performed.

## 3. Results

### 3.1. Augmentation effects of food additives on polysytrene consumption

To study the effects of food additives, mealworms and superworms were reared on PS with/without cinnamon, sucrose, and bran. Bran was the initial food source the worms were purchased with and was thus used as a control given previous reports of it increasing PS consumption [23]. Given the resistance of PS to natural decomposition, any loss of PS weight during the experiment is attributed to the consumption by worms. We found the addition of bran and sucrose to increase PS consumption by both superworms and mealworms. The addition of cinnamon increased PS consumption by superworms but had no significant effects on mealworms (Figure 2). Small amounts (25 mg) of bran were found to augment superworm PS consumption by 1.73 times, from 1.04 to 1.79 mg/g of worms/day (not statistically significant) and mealworm PS consumption by 1.53 times, from 1.40 to 2.14 mg/g of worms/day (not statistically significant). Similarly, the addition of sucrose increased superworm PS consumption by 1.83 times to 1.90 mg/g of worms/day (not statistically significant). The addition of sucrose increased mealworm PS consumption by 2.54 times to 3.55 mg/g of worms/day (P < 0.01). The mealworms significantly consumed more PS with sucrose than those cofed with bran (P < 0.05). Based on sucrose effects, mealworms significantly ate more PS than superworms (P < 0.05). With the exception of superworms fed on sucrose-PS recording a slight increase in weight of 1.79%. No significant worm weight change was found for the mealworms or superworms on PS-only diets over the four days (See Table S2 in the supplementary data).

**Figure 2.**
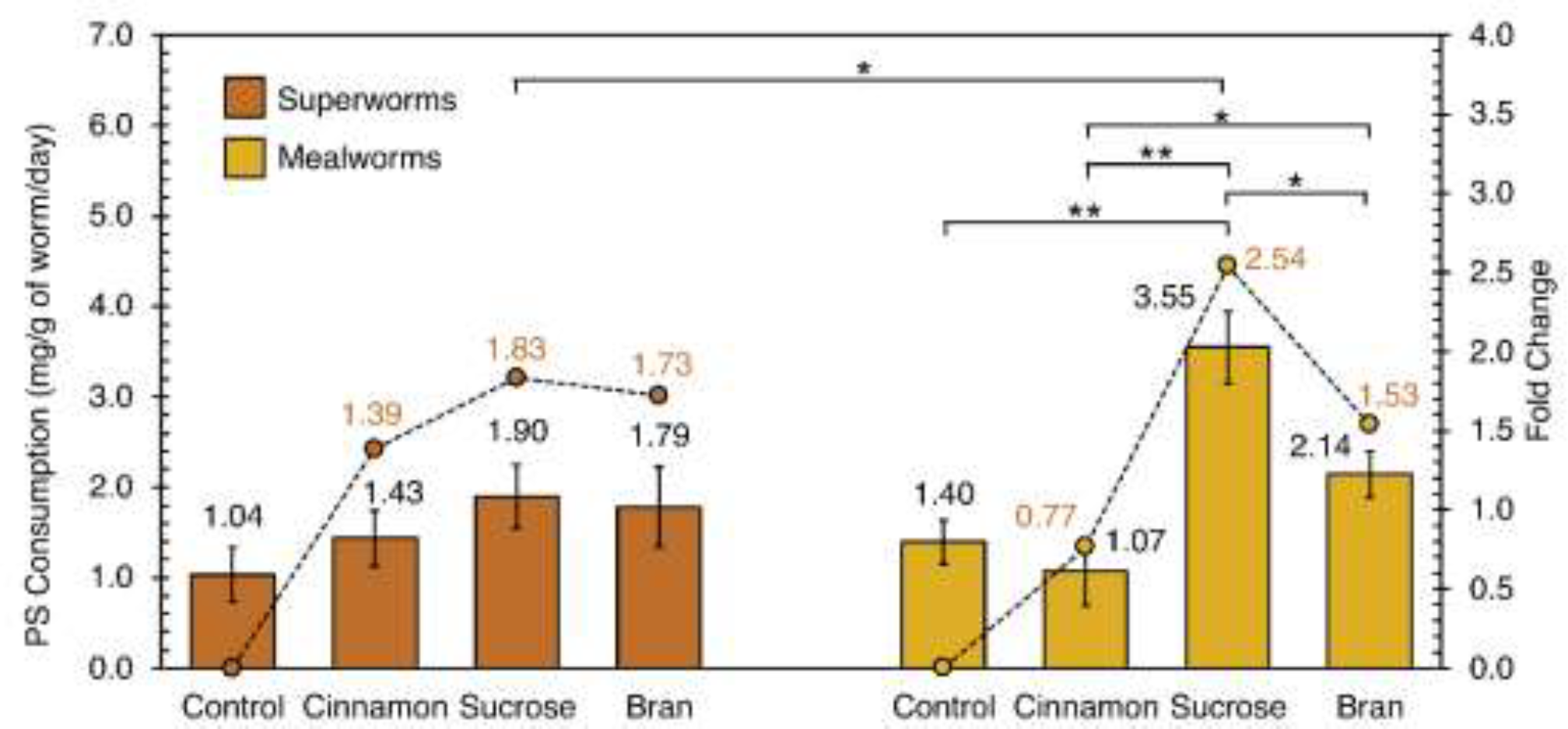
Rate of PS consumption (mg PS/g of worm/day) by superworms (left) and mealworms (right) with (cinnamon, sucrose and bran) and without (control) food additives. Additives were mixed with PS balls and sprayed with DI water to ensure adherence. Residual PS were weighed after four days. The bar chart shows the means of sextuplicates, with error bars representing the standard error. Statistical analysis was performed using two-tailed unequal variances T-test. *P < 0.05, **P < 0.01.

### 3.2. The effect of worm frass on cacti growth and rooting

Analysing cacti height growth, superworm frass media were found to support an average height gain of 0.50 cm that was not significantly different from those grown on tea leaves (average gain of 0.14 cm, see Figure 3). Mealworm frass media alone significantly impaired the growth of plants which lost an average height of 0.53 cm (P < 0.05), possibly due to water loss due to the poorer water retention properties of mealworm frass.

**Figure 3.**
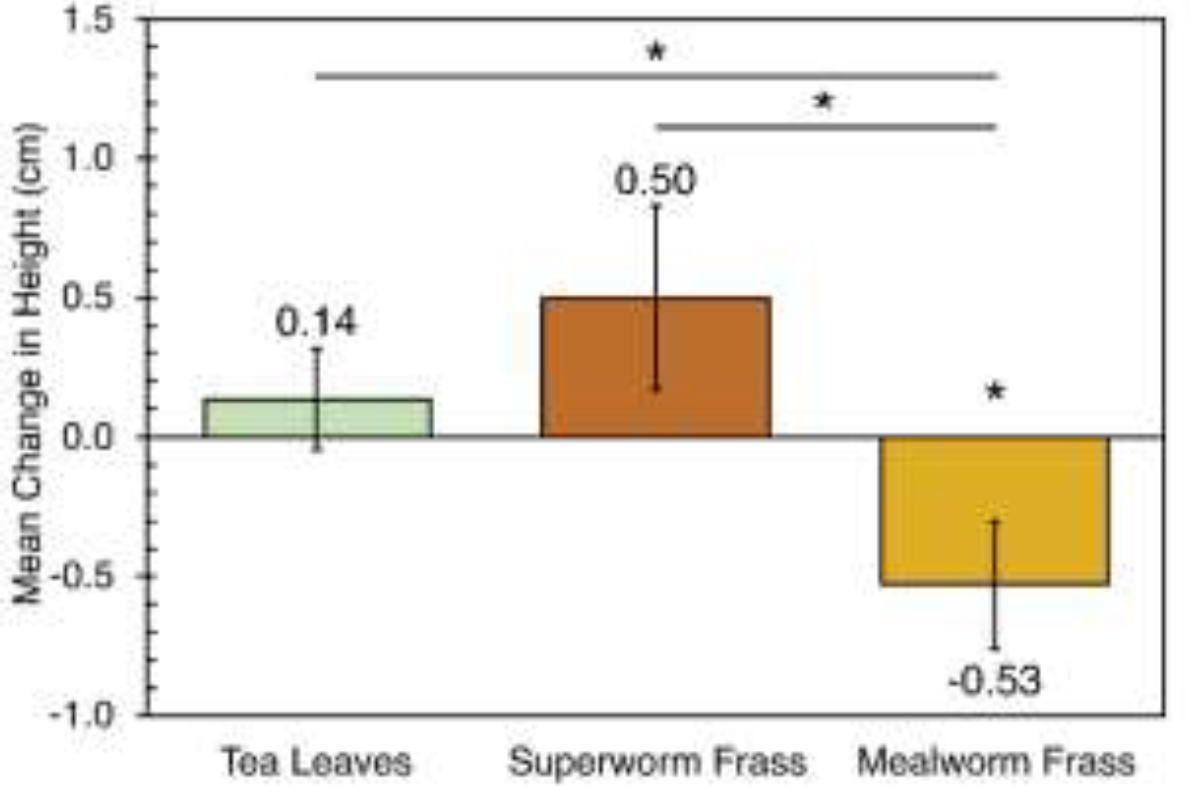
Effect of the three media on cacti growth. The error bars represent the standard error from 11 replicates. Statistical analysis was performed using two-tailed unequal variances T-test. *P < 0.05. The cacti were grown on the respective media for over a fortnight.

Frass from both superworms and mealworms fed solely on PS yielded a higher proportion of cacti rooting (below the soil line) than those grown on spent tea leaves (see Table 1). For frass from superworms fed on PS, nine cacti rooted (90.0%) compared to tea leaves with five cacti rooting (45.5 %, P < 0.05, see Table 1). Similarly, seven cacti grown on mealworm frass rooted (63.6%) compared to tea leaves (not statistically significant).

**Table 1.**
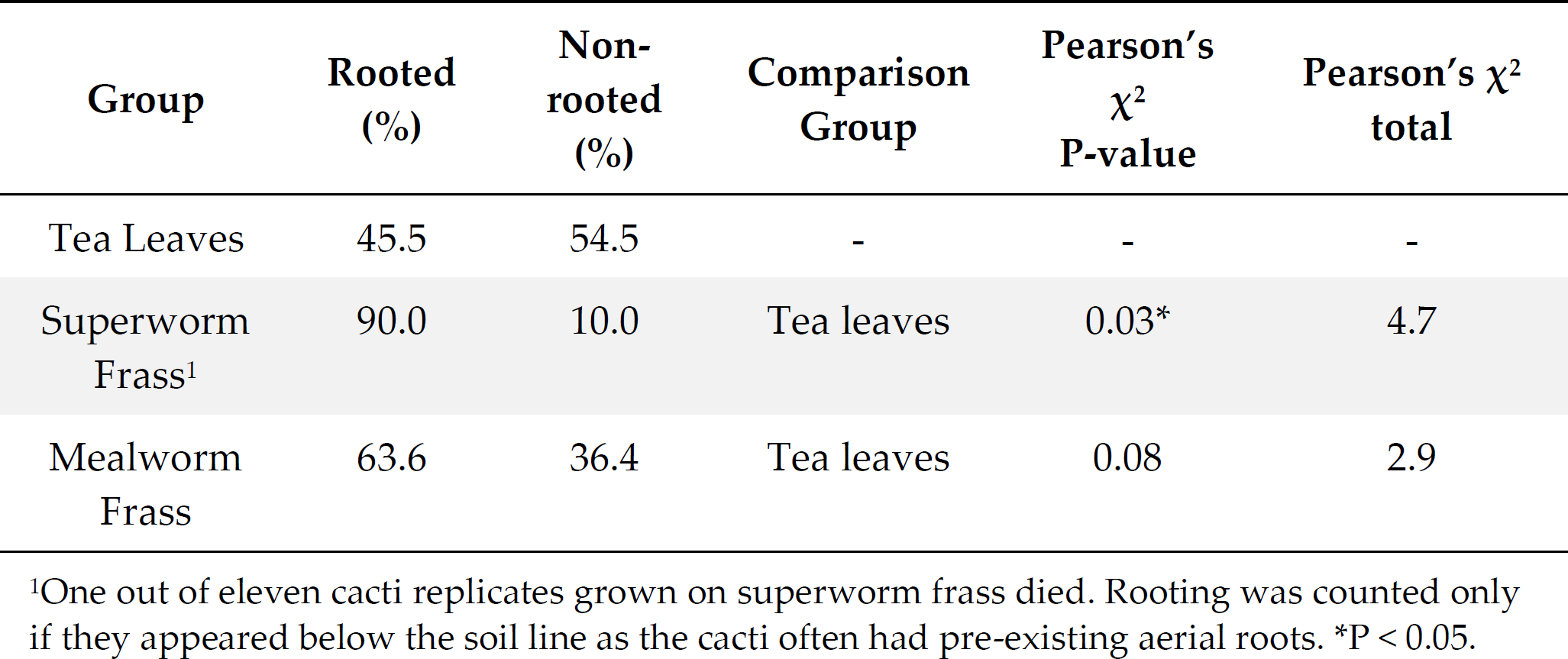
Effect of different media on number and proportions of rooting cacti.

### 3.3. GC-MS analysis of superworm frass

GC-MS detection of PS and by-products e.g., styrene in the superworm and mealworms did not show notable corresponding peaks (Figure 4) to those fed on bran. There were also no notable significant differences between the GC-MS analysis of the frass of the superworms fed on PS and that on bran (Figure S1). As a positive control, analysis of the PS balls alone showed peaks corresponding to styrene and molecules containing benzyl groups. Although without styrene and benzyl group peaks, the PS-fed worm frass samples had peaks corresponding to 9-oleamide (C18H35NO) fatty acid primary amides (FAPA) along with smaller peaks corresponding to mainly other FAPAs, short chain alkanes, alcohols, and cycloalkanes (Figure 4, Table S3).

**Figure 4.**
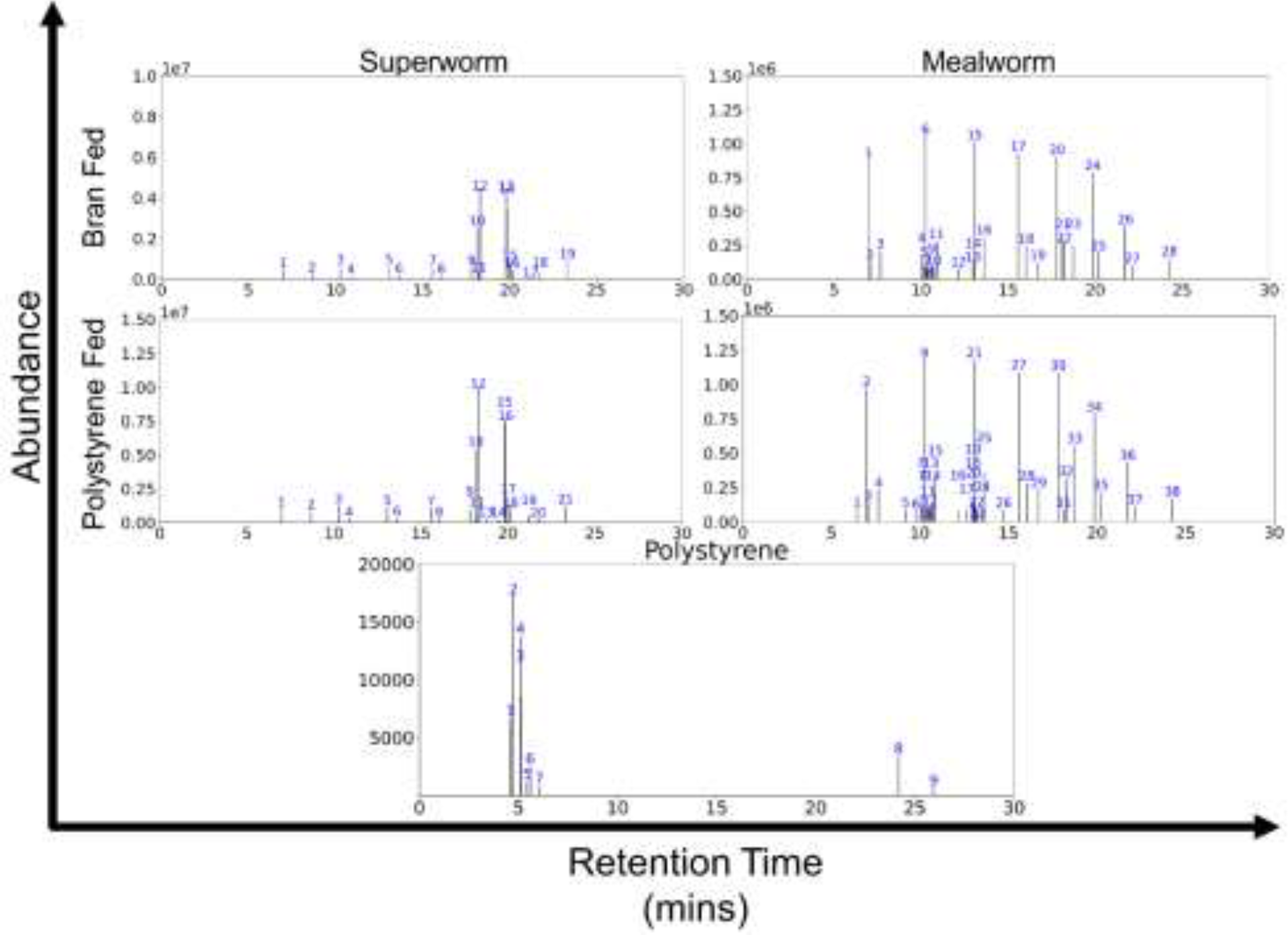
Representative GC-MS graphs from dissolved dead superworm (left) and mealworm (right), fed with bran (top) and polystyrene (bottom). Proposed chemicals corresponding to the identities of the different peaks is provided in Table S3.

## 4. Discussion

Studying the effects of food contamination of PS and the use of worm frass to support dragon fruit cacti growth, we find promise in using Superworms for a zero-waste circular economy.

Of the food additives, small additions of table sucrose (25 mg) were found to be the most effective, conferring mealworms greater consumption of the PS balls when compared to mealworms and superworms cofed on bran and sucrose, respectively. Mealworms and superworms fed with sucrose additives experienced the highest increase in PS consumption at 2.54 and 1.83-fold respectively. Bran, previously reported to double the rate of PS consumption [23], was also found to increase PS degradation of superworms by 1.73-fold and of mealworms by 1.53-fold. Our mealworms cofed with sucrose had a statistically significant increased consumption to mealworms cofed with bran. Since food is a major contaminant of PS waste, our findings suggest that food waste contamination to be an advantage for PS degradation by both mealworms and superworms. While there was weight loss of the worms over the 4 days of experimentation (see Table S1), it was not significant. We also did not notice hindering of mealworms life cycle from the plastic diet that was previously reported [26]. In general, we did not observe notable abnormalities during our worm breeding other than delayed pupation (which is beneficial for plastic degradation given that the larvae ate more than adult beetles) and have successfully bred subsequent generations of PS eating superworms and mealworms (unpublished data). All in all, our feeding experiment suggests that PS biodegradation using worms could be further optimized through the introduction of external factors.

### 4.1. Comparison of superworms to mealworms

There were no significant differences in PS consumption by worm weight between mealworms and superworms (for control conditions in Figure 2). While contrary to a recent report showing superiority of superworms over mealworms [20], the study calculated PS consumption per individual worm whereas we calculated by weight of the worm noting that a single superworm could be up to 20-folds heavier than a single mealworm. Given that actual counting of worms is not feasible when dealing with the tonnes of PS waste generated daily, we adopted weight measurements and recommend it for future scalability purposes.

### 4.2. Zero-waste circular economy

Mealworms and superworms are known fish [27] and poultry [12] feed. With the added plastic degrading ability [9,10,20,23,28] and their frass demonstrated as potential plant growth media, worms can address two major global problems of food production and PS waste with further research to ensure safety in the food chain to humans (see reviews on plasticizer accumulation in the food chain [29,30]). The notable advantage of mealworms and superworms over other insect larvae, such as black soldier flies used to deal with food waste [31] is that the mature darkling mealworm or superworm beetles have fused wings/elytra and do not fly, making their biocontainment significantly easier with minimal concern for their escape to cause local ecosystem problems.

The ability of mealworms in particular, to degrade plastic has been extensively studied [15,28]. Through GC-MS analysis, we did not detect styrene in the carcasses and frass of PS fed superworms nor notable differences from the bran-fed superworms. While more sensitive methods than GC-MS would be certainly necessary when seeking regulatory approval for food production, there is also much to be further investigated given that we used pure clean PS in our study. Given that real PS waste may include coloured or chemical additive-laced PS products, there is much to investigate before commercial implementation in real-life settings and introducing plastic eating worms into the food chain. Yet, our study here shows great promise of this approach and while likely not sensitive enough, a previous study utilising more sensitive gel permeation chromatography (GPC), Fourier-transform infrared spectroscopy (FTIR) and liquid-state ^1^H nuclear magnetic resonance (NMR) spectra characterised PS (of various foam material density) depolymerization and biodegradation by mealworms, finding polymer residues in their frass revealing evidence of partial depolymerization and oxidation [23].

Furthermore, previous reports [9,10,20,23,28] reported microplastics and plastic monomers in the frass or worms, yet given that PS degradation by mealworms and superworms increases progressively with time [9,20], there is promise to further optimise this process. We propose that the set-up of multi-layered chambers of worms where frass and small plastics falling to a lower chamber for further worm consumption with repeated iterations can allow for a complete removal of microplastics and partial depolymerization. It is also possible to utilize both mealworms and superworms simultaneously in such a setup as our other experiments (not shown) showed that apart from occasional cross-eating, they can be bred together, and even with black soldier fly larvae.

The dragon fruit cacti (*Hylocereus undatus*), an easy to grow indoor ornamental and food crop plant, was chosen for its hardiness to be an indicator plant in this study. Since superworm frass alone supported superior rooting and growth to the other media, it is recommended over mealworm frass (with lower proportion of rooting and loss in height, possibly due to its poorer ability to retain moisture). It is possible that short chain growth promoting alkene semiochemicals detected in the GC-MS analysis (e.g. Heptacosane, Nonadecane and Octadecane [32]), as well as chitin (reported to support rooting [33]) in the superworm frass augmented rooting. The researchers also noted that superworm frass was less pungent than the ammonia smelling mealworm frass making it possibly more suitable for indoor use. Nonetheless, the poor support of mealworm frass on cacti growth is unexpected given previous reports [15,16], although this could be due to the usage of 100% mealworm frass for our evaluation (as compared to it being used as a supplement) and the different nutritional requirements of the dragon fruit cacti that would require further study with other plants.

Given our findings that superworm frass is more suitable than mealworm frass, the combinatorial use of many plants reported to be able to clear up toxins from the environment [34] may be evaluated towards removal of any possible unwanted by-products from worm degradation that future research may uncover.

While there are exciting research based on enzymes isolated from bacteria present in plastic eating worms [35–38], complete degradation into harmless substances at industrial scale will require further research and engineering in the face of an urgently increasingly pressing problem of plastic waste made worse by lockdown measures during the COVID19 pandemic. In the meantime, the natural solution of worms can be investigated further for more immediate implementation, especially given their simultaneous roles as food, for urban farming in both fish/poultry feed and their frass for food crops. At the minimum, they reduce the PS waste to be sent for incineration. Worms are naturally more resistant to environmental factors compared to pure enzymes and can overcome obstacles for enzymes in plastic crystallinity or accessibility of the polymer chains, such that while protein engineering of such enzymes [39] are promising, there is still much to optimize before large scale implementation compared to worms.

The setups of both PS consumption by worms and frass-supported cacti growth were all performed indoors, demonstrating the possibility of worms as an environmentally friendly, widely implementable urban solution to plastic waste and food sustainability. Carried out within homes, this solution can significantly contribute to alleviating the global plastic waste and food production problem.

## 5. Conclusions

In conclusion, with evidence that food additives augment rather than antagonize PS degradation, and that the frass can be used to support food crop growth while the worms are themselves sources of human, poultry and fish feed, the answer in the worms is a very fitting scalable natural solution to both the plastic pollution and food (aquaculture and agriculture) production problems that could be implemented widely.

## Supplementary Materials

The following are available online at www.mdpi.com/xxx/s1,Table S1. Change in worm weight after four days with the different additives, Table S2. Effect of different media on mean height change of the cacti, Table S3. List of individual chemicals from a mass search of the peaks detected in GC-MS, Figure S1. GC-MS analysis of mealworm frass and superworm frass extracted from polystyrene fed worms.

## Author Contributions

Methodology, S.K.-E.G., S.-X.P. and Z.X.; formal analysis, S.-X.P., Z.X. and J.Y.Y.; investigation, S.-X.P. and Z.X.; writing – original draft preparation, S.K.-E.G.; writing – review and editing, S.K.-E.G., S.-X.P., Z.S.-L.H., J.Y.Y.; supervision, S.K.-E.G.; funding acquisition, S.K.-E.G.

## Institutional Review Board Statement

Not applicable.

## Informed Consent Statement

Not applicable.

## Data Availability Statement

Data presented in this study are available in the supplementary materials.

## Acknowledgments

The authors thank Ghin-Ray Goh, Benjamin Jun-Jie Aw, Elizabeth Yun-Ling Tay, Wen-Jie Tan, Xiaoyun Guan, Wai-Heng Lua, Darius Koh, Brennan Ang, and Wally Lam for assistance rendered in the husbandry of the worms and in parts of writing of the manuscript. We also thank Alfred Huan for access to the GC-MS.

