## Supplementary Data for "Method for Zero-Waste Circular Economy using worms for plastic agriculture: Augmenting polystyrene consumption and plant growth"

**Table S1.** Change in worm weight after four days with different additives.
Each set-up was performed in sextuplicate. Statistical analyses were performed using two-tailed Student's T-test. The average change in worm weight calculated in (g) and (%).

| Set-Up | | Initial Worm Weight (g) | Final Worm Weight (g) | Change in worm weight (g) | Average change in worm weight (g) | SEM | Average Change in worm weight (%) | t-test P value (two-Tailed) |
|---|---|---|---|---|---|---|---|---|
| **Superworms** | **Control** | 6.95 | 7.19 | 0.24 | <-0.01 | 0.08 | -0.04 | 0.97 |
| | | 7.42 | 7.46 | 0.04 | | | | |
| | | 7.46 | 7.55 | 0.09 | | | | |
| | | 10.40 | 10.38 | -0.02 | | | | |
| | | 10.70 | 10.67 | -0.03 | | | | |
| | | 10.33 | 9.99 | -0.34 | | | | |
| | **Cinnamon** | 6.98 | 6.93 | -0.05 | -0.11 | 0.09 | -1.24 | 0.27 |
| | | 7.08 | 7.11 | 0.03 | | | | |
| | | 6.77 | 6.97 | 0.20 | | | | |
| | | 10.66 | 10.48 | -0.18 | | | | |
| | | 10.40 | 10.14 | -0.26 | | | | |
| | | 10.33 | 9.94 | -0.39 | | | | |
| | **Sucrose** | 7.01 | 7.30 | 0.29 | 0.16 | 0.07 | 1.79 | 0.06 |
| | | 7.40 | 7.73 | 0.33 | | | | |
| | | 6.62 | 6.90 | 0.28 | | | | |
| | | 10.55 | 10.53 | -0.02 | | | | |
| | | 10.31 | 10.36 | 0.05 | | | | |
| | | 10.71 | 10.72 | 0.01 | | | | |
| | **Bran** | 6.57 | 6.67 | 0.10 | -0.06 | 0.08 | -0.69 | 0.50 |
| | | 7.26 | 7.13 | -0.13 | | | | |
| | | 7.05 | 7.30 | 0.25 | | | | |
| | | 10.10 | 9.99 | -0.11 | | | | |

| | | | | | | | | |
|---|---|---|---|---|---|---|---|---|
| | | 10.48 | 10.15 | -0.33 | | | | |
| | | 10.39 | 10.25 | -0.14 | | | | |
| **Mealworms** | **Control** | 6.79 | 6.36 | -0.43 | -0.24 | 0.22 | -2.85 | 0.32 |
| | | 6.29 | 5.37 | -0.92 | | | | |
| | | 6.44 | 5.65 | -0.79 | | | | |
| | | 10.49 | 10.58 | 0.09 | | | | |
| | | 10.46 | 10.70 | 0.24 | | | | |
| | | 10.76 | 11.11 | 0.35 | | | | |
| | **Cinnamon** | 6.22 | 5.74 | -0.48 | -0.20 | 0.16 | -2.36 | 0.27 |
| | | 6.23 | 5.89 | -0.34 | | | | |
| | | 6.45 | 5.89 | -0.56 | | | | |
| | | 10.54 | 10.63 | 0.09 | | | | |
| | | 10.44 | 10.90 | 0.46 | | | | |
| | | 10.45 | 10.09 | -0.36 | | | | |
| | **Sucrose** | 6.36 | 5.57 | -0.79 | -0.46 | 0.29 | -5.43 | 0.18 |
| | | 6.45 | 4.77 | -1.68 | | | | |
| | | 6.34 | 5.87 | -0.47 | | | | |
| | | 10.56 | 10.34 | -0.22 | | | | |
| | | 10.59 | 10.71 | 0.12 | | | | |
| | | 10.52 | 10.80 | 0.28 | | | | |
| | **Bran** | 6.40 | 5.84 | -0.56 | -0.17 | 0.12 | -1.98 | 0.23 |
| | | 6.11 | 5.76 | -0.35 | | | | |
| | | 6.59 | 6.20 | -0.39 | | | | |
| | | 10.69 | 10.84 | 0.15 | | | | |
| | | 10.75 | 10.89 | 0.14 | | | | |
| | | 10.50 | 10.50 | 0.00 | | | | |

**Table S2.** Effect of different media on mean change in height of the cacti.
Statistical analyses were performed using two-tailed T-tests. *P < 0.05, **P < 0.01.

| Plant Media | Initial Height (cm) | Final Height (cm) | Change in Height | | | | Two-Sample Unequal Variance T-test | |
|---|---|---|---|---|---|---|---|---|
| | | | Individual (cm) | Mean (cm) | SEM | Sig. (2-tailed) | Comparison Group | Sig. (2-tailed) |
| **Tea Leaves** | 9.4 | 9.5 | 0.1 | 0.14 | 0.18 | 0.83 | Superworm Frass | 0.36 |
| | 13.0 | 13.5 | 0.5 | | | | | |
| | 10.4 | 10.6 | 0.2 | | | | | |
| | 7.9 | 6.7 | 0.8 | | | | | |
| | 12.6 | 11.6 | -1.0 | | | | | |
| | 15.8 | 15.5 | -0.3 | | | | | |
| | 10.0 | 10.0 | 0.0 | | | | Mealworm Frass | 0.03* |
| | 10.9 | 10.6 | -0.3 | | | | | |
| | 8.2 | 8.0 | -0.2 | | | | | |
| | 9.0 | 10.1 | 1.1 | | | | | |
| | 20.1 | 20.7 | 0.6 | | | | | |
| **Superworm frass**[1] | 10.4 | 12.5 | 2.1 | 0.50 | 0.33 | 0.18 | Mealworm Frass | 0.02* |
| | 11.2 | 13.0 | 1.8 | | | | | |
| | 12.3 | 13.0 | 0.7 | | | | | |
| | 7.6 | 7.7 | 0.1 | | | | | |
| | 9.1 | 10.5 | 1.4 | | | | | |
| | 11.0 | 9.4 | -1.6 | | | | | |
| | 18.1 | 18.7 | 0.6 | | | | | |
| | 10.8 | 10.8 | 0.0 | | | | | |
| | 8.2 | 7.9 | -0.3 | | | | | |
| | 7.3 | 7.5 | 0.2 | | | | | |
| | 5.9 | Dead | | | | | | |
| **Mealworm frass** | 26.9 | 25.0 | -1.9 | -0.53 | 0.23 | 0.04* | - | - |
| | 20.9 | 19.1 | -1.8 | | | | | |
| | 13.1 | 13.9 | 0.8 | | | | | |

| | | |
|---|---|---|
| 13.0 | 12.8 | -0.2 |
| 13.6 | 13.5 | -0.1 |
| 8.1 | 7.6 | -0.5 |
| 7.5 | 7.3 | -0.2 |
| 6.1 | 5.4 | -0.7 |
| 7.7 | 7.1 | -0.6 |
| 9.9 | 9.6 | -0.3 |
| 11.9 | 11.6 | -0.3 |
| 8.1 | 7.6 | -0.5 |

[1] One out of eleven cacti replicates grown on superworm frass died

**Table S3.** List of individual chemicals from a mass search of the peaks detected in GC-MS.

| | PK | RT (mins) | Library search results |
|---|---|---|---|
| PS Balls | 1 | 4.63 | Benzene, ethyl |
| | 3 | 5.07 | 1,3,5,7-Cyclooctatetraene |
| | 4 | 5.1 | 3-Hexene, (E)- |
| | 7 | 6.05 | Benzene, propyl- |
| | 9 | 25.95 | Vanadium, (η7-cycloheptatrienylium)(η5-2,4-cyclopentadien-1-yl)- |
| Superworm fed with Polystyrene | 1 | 6.45 | Decane, 4-methyl- |
| | 2 | 6.99 | 3-Ethyl-3-methylheptane |
| | 3 | 7.07 | Decane, 3,7-dimethyl- |
| | 4 | 7.67 | Decane, 3,6-dimethyl- |
| | 5 | 9.19 | Pentadecane |
| | 6 | 9.75 | Tritetracontane |
| | 7 | 10.03 | 2,4-Dimethyldodecane |
| | 8 | 10.19 | Undecane, 2,9-dimethyl- |
| | 9 | 10.26 | Dodecane, 2,6,11-trimethyl- |
| | 10 | 10.38 | Octane, 5-ethyl-2-methyl- |
| | 11 | 10.45 | Dodecane, 1-iodo- |
| | 12 | 10.56 | Hexadecane |
| | 13 | 10.63 | Dichloroacetic acid, 6-ethyl-3-octyl ester |
| | 14 | 10.74 | Hexane, 2,3,4-trimethyl- |
| | 15 | 10.89 | Dodecane, 2,6,11-trimethyl- |
| | 16 | 12.16 | Heneicosane |
| | 17 | 12.65 | Hexadecane |
| | 18 | 12.97 | Tetracosane |
| | 19 | 13.01 | Hexadecane |
| | 20 | 13.09 | Octacosane |
| | 21 | 13.20 | Hexadecane |
| | 22 | 13.30 | Phenol, 2,4-bis(1,1-dimethylethyl)- |
| | 23 | 13.63 | Octacosane |
| | 24 | 14.74 | Heneicosane |
| | 25 | 15.59 | Tridecane, 5-propyl- |
| | 26 | 16.05 | Heneicosane |
| | 27 | 16.72 | Hexadecanal |
| | 28 | 17.83 | Heneicosane |
| | 29 | 18.13 | Octacosane |
| | 30 | 18.24 | Heptadecane, 9-octyl- |
| | 31 | 18.75 | Oxirane, hexadecyl- |
| | 32 | 19.85 | Octacosane |
| | 33 | 20.22 | Octacosane |
| | 34 | 21.75 | Heptacosane |
| | 35 | 22.16 | Hexadecane, 2-methyl- |
| | 36 | 24.27 | Heptacosane |
| Mealworm fed with Polystyrene | 1 | 6.99 | 3-Ethyl-3-methylheptane |
| | 2 | 7.67 | Undecane, 4,7-dimethyl- |
| | 3 | 8.70 | Octanoic Acid |
| | 4 | 10.26 | Dodecane, 2,6,11-trimethyl- |
| | 5 | 10.89 | Dodecane, 2,6,11-trimethyl- |
| | 6 | 13.09 | Tridecane, 1-iodo- |

| | | | |
|---|---|---|---|
| | 7 | 13.63 | Pentadecane |
| | 8 | 15.58 | Pentadecane |
| | 9 | 16.05 | Octadecane |
| | 10 | 17.83 | Heptacosane |
| | 11 | 18.17 | n-Hexadecanoic acid |
| | 12 | 18.24 | Pentacosane |
| | 13 | 18.32 | Octanoic acid, decyl ester |
| | 14 | 18.75 | Hexadecanal |
| | 15 | 19.46 | Heneicosane |
| | 16 | 19.83 | 9,12-Octadecadienoic acid (Z,Z)- |
| | 17 | 19.86 | 9-Octadecenoic acid, (E)- |
| | 18 | 20.04 | Octadecanoic acid |
| | 19 | 20.16 | Cyclododecane |
| | 20 | 21.23 | Hexadecane |
| | 21 | 21.75 | Heptacosane |
| | 22 | 23.33 | Pentadecane, 8-hexyl- |
| Superworm fed with bran | 1 | 6.99 | Decane, 2,3,7-trimethyl- |
| | 2 | 7.08 | Octane, 5-ethyl-2-methyl- |
| | 3 | 7.67 | Nonane, 2,6-dimethyl- |
| | 4 | 10.03 | Oxalic acid, 6-ethyloct-3-yl isohexyl ester |
| | 5 | 10.19 | Hexane, 3,3-dimethyl- |
| | 6 | 10.26 | Sulfurous acid, 2-ethylhexyl hexyl ester |
| | 7 | 10.38 | Decane, 2,3,7-trimethyl- |
| | 8 | 10.46 | Tridecane |
| | 9 | 10.63 | Decane, 1,1'-oxybis- |
| | 10 | 10.75 | Cyclopentane, propyl- |
| | 11 | 10.89 | Dodecane, 2,6,11-trimethyl- |
| | 12 | 12.16 | Heptadecane |
| | 13 | 12.98 | Decane, 2,4,6-trimethyl- |
| | 14 | 13.01 | Hexadecane |
| | 15 | 13.09 | Sulfurous acid, 2-ethylhexyl isohexyl ester |
| | 16 | 13.64 | Eicosane |
| | 17 | 15.59 | Tridecane, 5-propyl- |
| | 18 | 16.06 | 2-Bromo dodecane |
| | 19 | 16.73 | Tetradecanal |
| | 20 | 17.83 | Heneicosane |
| | 21 | 18.15 | n-Hexadecanoic acid |
| | 22 | 18.24 | Heneicosane |
| | 23 | 18.76 | Oxirane, hexadecyl- |
| | 24 | 19.86 | Heptacosane |
| | 25 | 20.22 | Heptadecane, 9-octyl- |
| | 26 | 21.76 | Tetratriacontane |
| | 27 | 22.16 | Tetratriacontane |
| | 28 | 24.28 | Heptacosane |
| Mealworm fed with bran | 1 | 6.99 | Dodecane |
| | 2 | 8.66 | Octanoic Acid |
| | 3 | 10.27 | Sulfurous acid, 2-ethylhexyl hexyl ester |
| | 4 | 10.89 | Dodecane, 2,6,11-trimethyl- |
| | 5 | 13.09 | Tridecane, 1-iodo- |
| | 6 | 13.64 | Tridecane, 1-iodo- |
| | 7 | 15.59 | Heneicosane |

| | | | |
|---|---|---|---|
| | 8 | 16.06 | Octacosane |
| | 9 | 17.83 | Heptacosane |
| | 10 | 18.16 | n-Hexadecanoic acid |
| | 11 | 18.24 | Octacosane |
| | 12 | 18.32 | Octanoic acid, dodecyl ester |
| | 13 | 19.82 | 9,12-Octadecadienoic acid (Z,Z)- |
| | 14 | 19.86 | 9-Octadecenoic acid, (E)- |
| | 15 | 20.04 | Octadecanoic acid |
| | 16 | 20.17 | Cyclododecane |
| | 17 | 21.24 | Eicosane |
| | 18 | 21.76 | Eicosane |
| | 19 | 23.35 | Eicosane, 10-methyl- |
| | 1 | 3.80 | 2,4-Dimethyl-1-heptene |
| | 2 | 6.31 | Octane, 3,4,5,6-tetramethyl- |
| | 3 | 6.44 | Octane, 3-ethyl- |
| | 4 | 6.99 | 3-Ethyl-3-methylheptane |
| | 5 | 7.08 | 2-Bromononane |
| | 6 | 7.33 | 4-Methyl-2-heptene |
| | 7 | 7.39 | 1-Decene, 8-methyl- |
| | 8 | 7.67 | Decane, 3,6-dimethyl- |
| | 9 | 8.71 | 1,4-Dioxane-2,5-dione, 3,6-dimethyl-, (3S-cis)- |
| | 10 | 9.19 | Decane, 2,4,6-trimethyl- |
| | 11 | 9.75 | Tritetracontane |
| | 12 | 10.03 | Oxalic acid, bis(6-ethyloct-3-yl) ester |
| | 13 | 10.19 | Nonane, 3-methyl-5-propyl- |
| | 14 | 10.26 | Dodecane, 2,6,11-trimethyl- |
| | 15 | 10.38 | Octane, 2,4,6-trimethyl- |
| | 16 | 10.45 | Dodecane, 4,6-dimethyl- |
| | 17 | 10.63 | Sulfurous acid, isohexyl 2-pentyl ester |
| | 18 | 10.74 | 1-Hexene, 3,3-dimethyl- |
| Frass from mealworms fed on polystyrene | 19 | 10.89 | Decane, 6-ethyl-2-methyl- |
| | 20 | 12.16 | Heneicosane |
| | 21 | 12.65 | Hexadecane |
| | 22 | 12.97 | Heptadecane |
| | 23 | 13.01 | Heptadecane |
| | 24 | 13.09 | Tridecane, 1-iodo- |
| | 25 | 13.31 | Phenol, 2,4-bis(1,1-dimethylethyl)- |
| | 26 | 13.55 | 2-Bromotetradecane |
| | 27 | 13.63 | Tridecane, 1-iodo- |
| | 28 | 14.74 | Heneicosane |
| | 29 | 15.59 | Heneicosane |
| | 30 | 16.06 | Pentacosane |
| | 31 | 16.72 | Hexadecanal |
| | 32 | 17.83 | Eicosane, 2-methyl- |
| | 33 | 18.13 | Nonahexacontanoic acid |
| | 34 | 18.24 | Octadecane, 1-iodo- |
| | 35 | 18.75 | Oxirane, heptadecyl- |
| | 36 | 19.85 | Decane, 2,9-dimethyl- |
| | 37 | 20.22 | Tetracosane |
| | 38 | 21.75 | Heptacosane |
| | 39 | 22.16 | Tetratriacontane |

| | | | |
|---|---|---|---|
| | 40 | 22.95 | Benzonitrile, m-phenethyl- |
| | 41 | 23.34 | Pentacosane |
| | 42 | 24.27 | Octacosane |
| Frass from superworms fed on polystyrene | 1 | 3.80 | 2,4-Dimethyl-1-heptene |
| | 2 | 6.31 | Octane, 3,4,5,6-tetramethyl- |
| | 3 | 6.44 | Decane, 4-methyl- |
| | 4 | 6.89 | Sulfurous acid, hexyl pentyl ester |
| | 5 | 6.99 | 3-Ethyl-3-methylheptane |
| | 6 | 7.07 | Decane, 4-ethyl- |
| | 7 | 7.33 | Nitric acid, nonyl ester |
| | 8 | 7.39 | Cyclopropane, 1-butyl-2-pentyl-, trans- |
| | 9 | 7.67 | Decane, 3,6-dimethyl- |
| | 10 | 9.19 | Tetradecane |
| | 11 | 9.75 | Methoxyacetic acid, 2-tetradecyl ester |
| | 12 | 10.03 | Decane, 1-iodo- |
| | 13 | 10.19 | Heptadecane |
| | 14 | 10.26 | Dodecane, 2,6,11-trimethyl- |
| | 15 | 10.38 | Dodecane, 2,6,11-trimethyl- |
| | 16 | 10.45 | Dodecane |
| | 17 | 10.56 | Tridecane, 1-iodo- |
| | 18 | 10.63 | Sulfurous acid, isohexyl 2-pentyl ester |
| | 19 | 10.74 | Cyanamide, dibutyl- |
| | 20 | 10.89 | Dodecane, 2,6,11-trimethyl- |
| | 21 | 12.16 | Heptadecane |
| | 22 | 12.65 | Hexadecane |
| | 23 | 12.98 | Dodecane, 3-methyl- |
| | 24 | 13.01 | Dodecane, 4,6-dimethyl- |
| | 25 | 13.09 | Hexadecane |
| | 26 | 13.31 | Phenol, 2,4-bis(1,1-dimethylethyl)- |
| | 27 | 13.55 | Decane, 3,6-dimethyl- |
| | 28 | 13.63 | Octacosane |
| | 29 | 14.74 | Heneicosane |
| | 30 | 15.59 | Heneicosane |
| | 31 | 16.06 | Pentadecane |
| | 32 | 16.72 | Tetradecanal |
| | 33 | 17.06 | Heneicosane |
| | 34 | 17.83 | Octadecane, 1-iodo- |
| | 35 | 18.13 | Methoxyacetic acid, 2-tridecyl ester |
| | 36 | 18.24 | Heneicosane |
| | 37 | 18.75 | Oxirane, hexadecyl- |
| | 38 | 19.85 | Octadecane |
| | 39 | 20.22 | Tetratriacontane |
| | 40 | 21.69 | Eicosane |
| | 41 | 21.76 | Octadecane, 1-iodo- |
| | 42 | 22.16 | Eicosane |
| | 43 | 24.27 | Heptacosane |

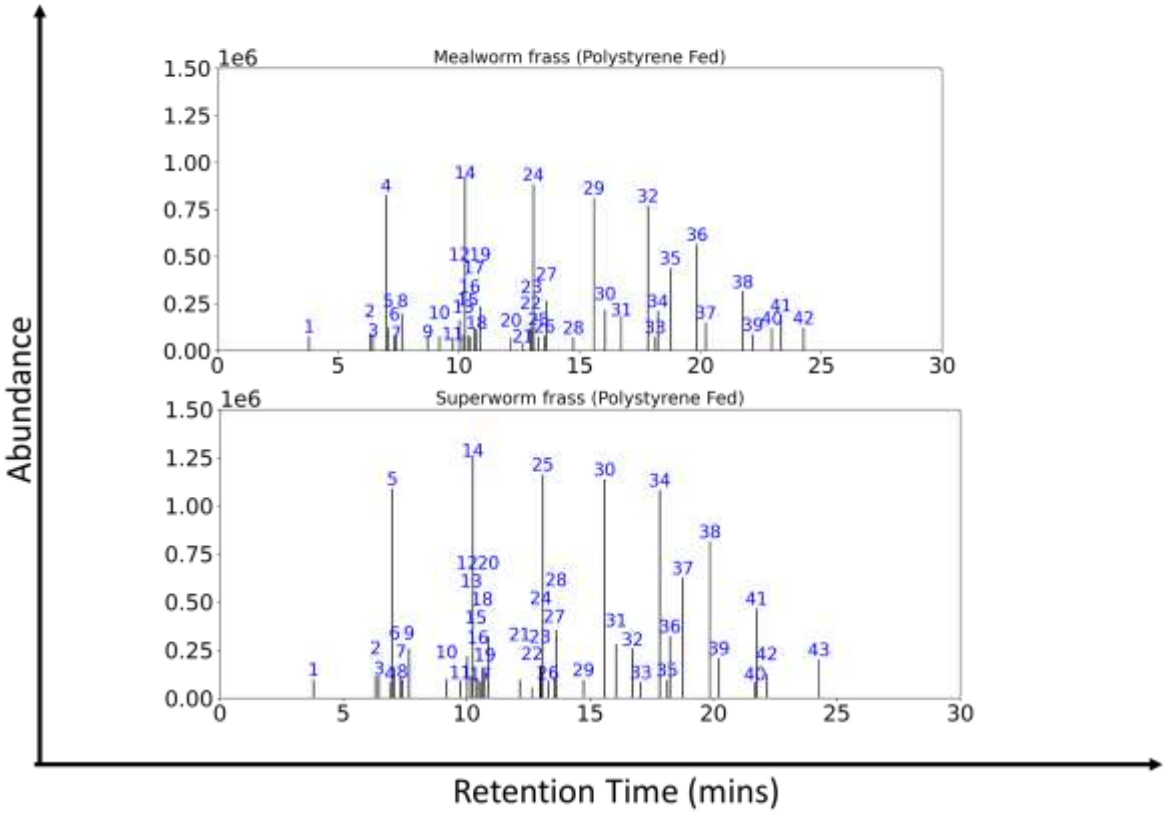


**Figure S1.** GCMS analysis of mealworm frass and superworm frass extracted from polystyrene fed worms
